# Mapping the tumor stress network reveals dynamic shifts in the stromal oxidative stress response

**DOI:** 10.1101/2023.09.29.560126

**Authors:** Chen Lior, Debra Barki, Christine A Iacobuzio-Donahue, David Kelsen, Ruth Scherz-Shouval

## Abstract

The tumor microenvironment (TME) is a challenging environment where cells must cope with stressful conditions such as fluctuating pH levels, hypoxia, and free radicals. In response, stress pathways are activated, which can both promote and inhibit tumorigenesis. In this study, we set out to characterize the stress response landscape across four carcinomas: breast, pancreas, ovary, and prostate tumors, focusing on five pathways: Heat shock response, oxidative stress response, unfolded protein response, hypoxia stress response, and DNA damage response. Using a combination of experimental and computational methods, we create an atlas of the stress response landscape across various types of carcinomas. We find that stress responses are heterogeneously activated in the TME, and highly activated near cancer cells. Focusing on the non-immune stroma we find, across tumor types, that NRF2 and the oxidative stress response are distinctly activated in immune-regulatory cancer-associated fibroblasts and in a unique subset of cancer associated pericytes. Our study thus provides an interactome of stress responses in cancer, offering new ways to intersect survival pathways within the tumor, and advance cancer therapy.

## Introduction

Cancer development and progression is a complex process involving not only malignant cells but also the surrounding tumor microenvironment (TME), comprising various non-malignant cells such as fibroblasts, pericytes, and immune cells. These cells face a stressful environment due to nutrient scarcity, hypoxia, fluctuating pH levels, and demands for rapid protein translation, necessitating the activation of survival pathways (Akman, 2021; Seebacher et al, 2021; Leprivier et al, 2015).

Within the tumor microenvironment, cancer-associated fibroblasts (CAFs) and pericytes are among the most abundant cell types in various carcinomas, and they contribute significantly to cancer progression (Ping et al, 2021; Sun et al, 2021). CAFs, a heterogeneous population originating from various sources, are generally divided into three - myofibroblastic CAFs (myCAFs), immune-regulatory CAFs (iCAFs), and antigen-presenting CAFs (apCAFs)(Sahai *et al*, 2020; Ping *et al*, 2021; Santi *et al*, 2018; Ganguly *et al*, 2020; Liu *et al*, 2019; Chen *et al*, 2021b). They interact with other cells of the TME, such as immune cells, facilitating a pro-tumorigenic environment (Liu *et al*, 2019; Lavie *et al*, 2022; Elyada *et al*, 2019; Arpinati & Scherz-Shouval, 2023; Mun *et al*, 2022). Pericytes, mural cells of blood vessels, are involved in tumor angiogenesis and metastasis, regulating vascular stability, and enhancing tumor cell intravasation when dysfunctional (Armulik *et al*, 2011; Sun *et al*, 2021). Although these cells evidently play a significant role in tumor progression, our understanding of their transformation into cancer-associated states and their mechanisms of influence remains a topic of active research (Ping *et al*, 2021; Ganguly *et al*, 2020; Liu *et al*, 2019; Chen *et al*, 2021b; Kharaishvili *et al*, 2014).

Cellular stress responses, including the unfolded protein response (UPR) (Hetz, 2012), heat shock response (HSR) (Richter *et al*, 2010), oxidative stress response, (OSR) (Sies & Jones, 2020) hypoxia stress response, (HySR) (Semenza, 2014), and the DNA damage response (DDR) (Lord & Ashworth, 2012), help maintain cellular homeostasis and survival under adverse conditions. In the context of cancer, these pathways have a dual role: they can promote survival and thus facilitate tumorigenesis, however chronic activation of them can lead to cell death, potentially inhibiting tumor growth (Siwecka *et al*, 2019). For example, both the HSR and the UPR can promote cancer cell survival by stabilizing protein folding and reducing protein aggregation (Li *et al*, 2011; Madden *et al*, 2019; Cyran & Zhitkovich, 2022); hypoxic conditions were shown to be beneficial for the tumor by promoting vascularization and angiogenesis (Li *et al*, 2021; Krock *et al*, 2011; Sebestyén *et al*, 2021), and mutations in DNA damage response genes, such as *BRCA1* and *BRCA2*, can result in genome instability and an increased risk of developing breast and ovarian cancers (Roy *et al*, 2012). Understanding these cellular stress responses within the TME is crucial for developing novel cancer therapies, highlighting the need for further research into the specific mechanisms and signaling pathways involved, their interactions, and their potential as therapeutic targets.

In recent years there have been numerous studies exploring the potential roles of stress responses in various cellular components of the TME (Varone *et al*, 2021; Grunberg *et al*, 2020; Zhang *et al*, 2013; Ramirez *et al*, 2020; Nguyen *et al*, 2018; Chen & Cubillos-Ruiz, 2021; Miles *et al*, 2019). Work by us and others described the importance of different stress responses in CAFs, and highlighted non­cell-autonomous roles for stress responses (Martinez-Outschoorn *et al*, 2010; Verginadis *et al*, 2022; Matsuzaki *et al*, 2015; Chan *et al*, 2017; Scherz-Shouval *et al*, 2014; Grunberg *et al*, 2021; Levi-Galibov *et al*, 2020; Shaashua *et al*, 2022). However, these studies have largely focused on individual stress responses. We lack a comprehensive description of the stress network. Given the diversity and intercommunication among cells within the TME, a nuanced, cell-specific understanding of how stress responses influence each cellular compartment and interact with each other is pivotal.

In this study, we took a holistic approach and examined the network of stress responses in the tumor and its microenvironment. We utilized multiplexed immunofluorescence (MxIF) staining of human patient samples to characterize activation patterns of stress responses across carcinomas in four different organs - pancreas, breast, ovary, and prostate. We found a gradient of activation, whereby stromal cells located closer to the cancer cells exhibit higher stress response activation levels. Analysis of patient-derived single-cell RNA-sequencing (scRNA-seq) data allowed us to create a cell- and organ-specific atlas of the stress response landscape across various types of carcinomas. Through our analysis, we discerned distinct subpopulations of fibroblasts and pericytes that exhibit a clear association with cellular stress, in particular oxidative stress, orchestrated by the transcription factor NRF2. This comprehensive map and the identified molecular interactions pave the way to elucidate the contribution of stress responses in the tumor microenvironment in a cell-specific manner. Moreover, it offers insights into how the stress response landscape might influence tumor progression and disease outcome.

## Results

### Stress responses are heterogeneously activated in the stroma, and their activation increases with proximity to cancer cells

To map the stress response network in the TME, we monitored the activation status in human tumors of transcription factors driving three major stress response pathways. We stained human tumor microarrays (TMAs) derived from four tumor types - breast, pancreas, ovary, and prostate - with antibodies for three stress transcription factors (TFs): NRF2 (Nuclear factor erythroid 2- related factor 2), a key factor of the oxidative stress response (OSR); ATF4 (Activating transcription factor 4), a master regulator of the unfolded protein stress response (UPR) and the integrated stress response; and HSF1 (Heat shock factor 1), which orchestrates the heat shock stress response (HSR) (Figure 1A). These TFs translocate to the nucleus upon activation and therefore their localization can be used as a proxy to monitor activation (Shaashua *et al*, 2022). Non-immune stromal cells were identified by negative staining for CD45 (immune cells) and CK (epithelial cells). Across all tumors, stress TFs were more strongly activated in cancer cells compared to non-malignant cells in the TME, as expected(Chen & Xie, 2018). Nevertheless, we observed marked activation of stress TFs in stromal cells, which appeared to be spatially heterogeneous (Figure 1A). To assess this spatial diversity, we calculated the distance between each non-immune stromal cell and its nearest cancer cell and examined whether this distance differs between stressed and unstressed cells (defined as cells that stained positively for one of the stress TFs, see Methods). We found that, across tumor types, stressed stromal cells are localized significantly closer to the cancer cells compared to unstressed stromal cells (Figure 1B). Analyzing each stress pathway separately, we observed a negative correlation between the staining intensity of each stress marker and the distance of the stromal cell to the nearest cancer cell (Figure 1C). This was true across all tumor types, yet it was most pronounced in ovarian tumors and least evident in pancreatic tumors, suggesting not only that the spatial heterogeneity of stress responses varies among tumor types but also that cancer cells might transfer, confer, or induce stress to the stroma.

**Figure 1.**
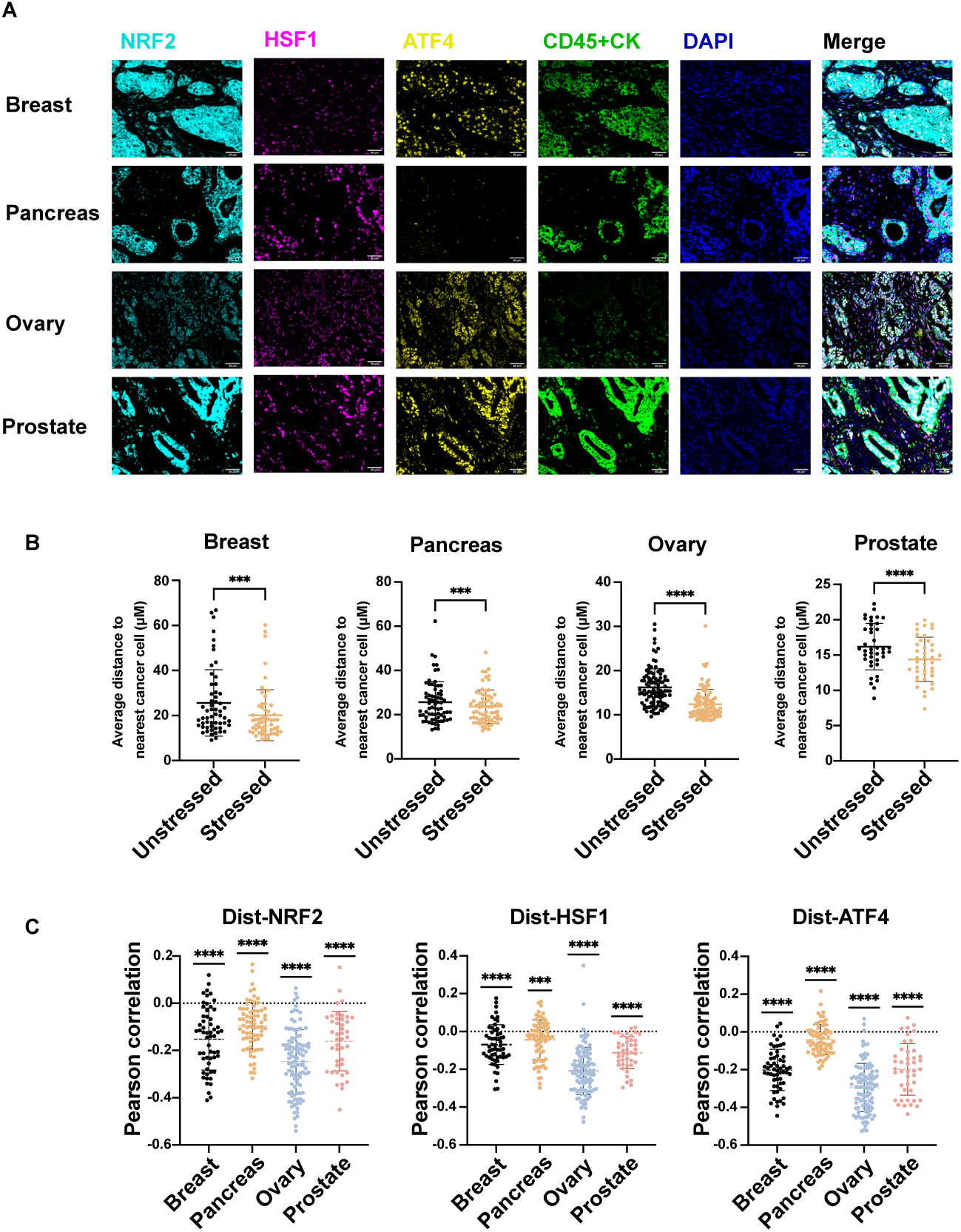
Stromal stress response activation increases with proximity to cancer cells. Formalin-fixed paraffin-embedded (FFPE) tumor microarrays (TMAs) of breast (N = 57), pancreas (N = 71), ovary (N = 102) and prostate (N = 43) cancer patients were stained by MxIF for the indicated proteins. DAPI was used to stain nuclei. **(A)** Representative images are shown. Scale bar = 50 μm. **(B)** Images were analyzed using QuPath software, CD45^-^ CK^-^ cells were defined as non-immune stromal cells, and stratified to stressed and unstressed cells based on staining for either NRF2, HSF1, or ATF4. The distance of each non-immune stromal cell to its nearest cancer cell was calculated and averaged. **(C)** For each patient, the Pearson correlation coefficient between the intensity of the indicated protein of non-immune stromal cells and the distance to the nearest cancer cell was calculated. P-Values were calculated using the Student t-test (**B**-paired t-test, **C**-one sample t-test (μ=0)).

### Transcriptomic analysis uncovers universal and organ-specific non-cell autonomous stress response activation patterns

To gain a better understanding of the stress response landscape in the TME, we evaluated the transcriptional patterns of the different stress responses in the TME using publicly available scRNA-seq data of the four tumor types (breast, pancreas, ovary, prostate) (Pal *et al*, 2021; Wu *et al*, 2021; Chen *et al*, 2021a; Werba *et al*, 2023; Steele *et al*, 2020; Geistlinger *et al*, 2021; Zhang *et al*, 2022; Olbrecht *et al*, 2021; Peng *et al*, 2019) (Figure 2A-D). To the three stress responses evaluated by immunostaining (OSR, HSR and UPR), we added the cellular response to hypoxia (HySR) and the DNA damage response (DDR). We generated a score for each stress response based on the average expression levels of a signature of target genes (50-250 genes each, see Methods, Supplementary Table 1). For each patient, we determined the mean expression for each of the five stress scores and then compared the scores across different cell types (Figure 2E-I; Supplementary Figure 1A- E). We found distinct patterns of expression in the different cell types, which were largely shared across tumor types. OSR scores were highest in epithelial, myeloid, pericytes and fibroblasts (Figure 2E; Supplementary Figure 1A), while the hypoxia stress response was highest in endothelial cells in breast, pancreas, and prostate tumors, and was among the highest in the endothelial cells of ovarian tumors (Figure 2F; Supplementary Figure 1B). HSR scores were divergent in their distribution across cell types in the different tumors (Figure 2G; Supplementary Figure 1C), and B cells expressed high UPR scores in all tumors, potentially due to the protein folding stress associated with the requirement to translate and sustain a high level of antibodies (Jiang *et al*, 2021) (Figure 2H; Supplementary Figure 1D). T, B, and myeloid cells expressed high DNA damage response scores in all tumors (Figure 2I; Supplementary Figure 1E). To assess not only the individual stress responses but their potential co-regulation, we calculated correlations between the stress scores of each cell type across patients (Figure 3A-D). While different organs have different stress networks, we observe some shared characteristics: In breast, pancreas, and ovarian tumors there is a strong co-regulation of non-immune-stromal HySR - (fibroblasts, endothelial cells and/or pericytes; Figure 3A-C), and the DDR is co-regulated in different immune cell types in breast, pancreas, and prostate tumors, indicating that these stresses are experienced similarly in those cell types. The HSR appears to be the most global stress response in prostate tumors, while in pancreatic tumors it appears to be the DDR, and in ovarian tumors the UPR, indicated by a co-regulation pattern across most cell types (Figure 3A-D). In breast tumors, both HSR and OSR are strongly correlated in the non-immune stroma. Taken together, the imaging and scRNA-seq analysis suggest an inter-cellular communication network of stress responses in the TME, with global as well as organ-specific characteristics.

**Figure 2.**
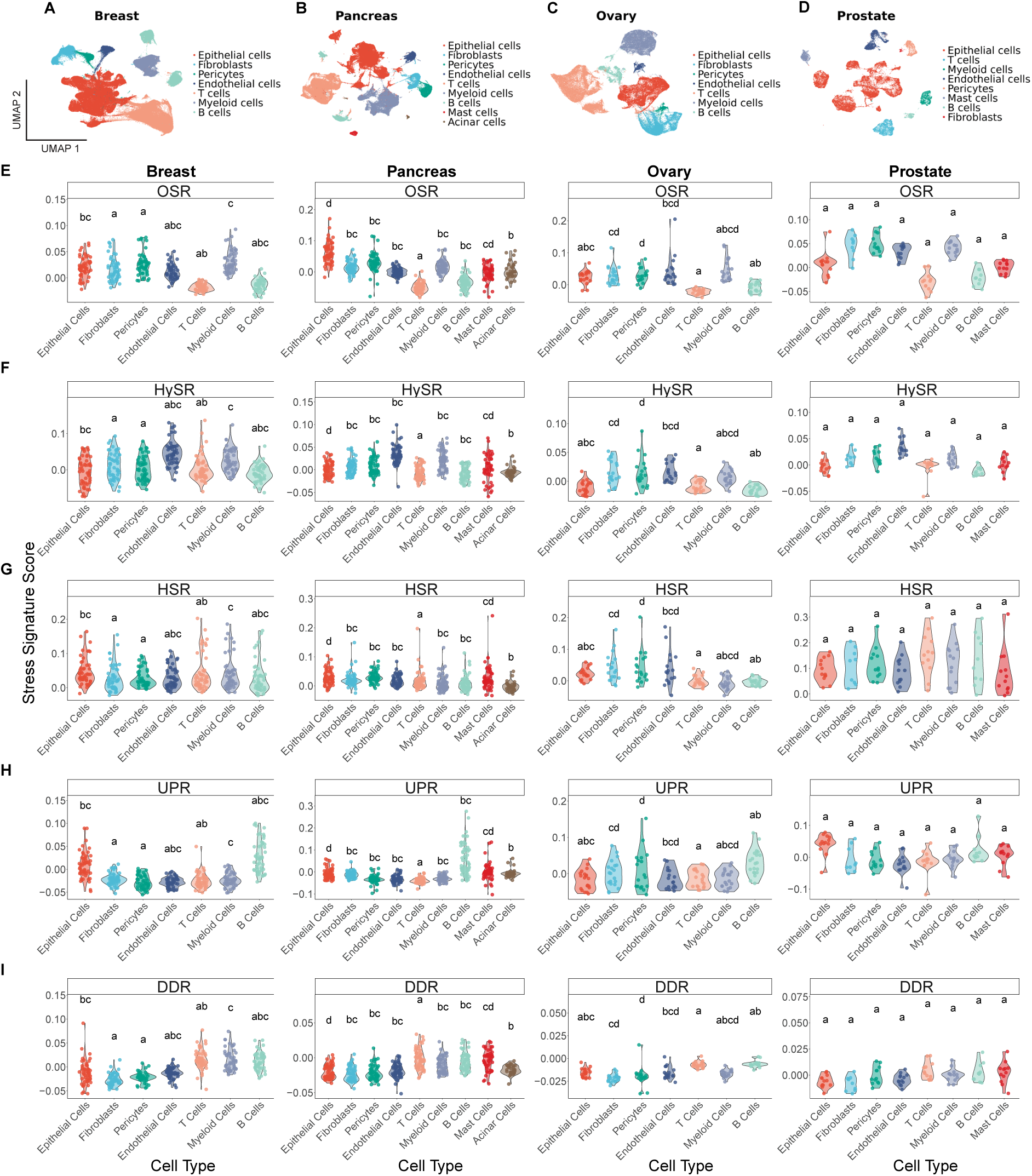
scRNA-seq data analysis uncovers shared and unique stress response patterns across organs. scRNA-seq data from human breast, pancreas, ovary, and prostate tumors was eanalyzed using the Seurat R toolkit. **(A-D)** UMAP plots of 265,034 cells from 51 breast cancer *atients (Pal *et al*, 2021; Wu *et al*, 2021) **(A);** 199,938 cells from 59 pancreatic cancer patients Werba *et al*, 2023; Steele *et al*, 2020; Peng *et al*, 2019) **(B);** 84,369 cells from 20 ovarian cancer * atients (Geistlinger *et al*, 2021; Zhang *et al*, 2022; Olbrecht *et al*, 2021) **(C);** and 32,823 cells from 3 prostate cancer patients (Chen *et al*, 2021a) **(D).** UMAPs are colored by cell type, defined by differential gene expression and canonical cell type markers. **(E-I)** Quantification of stress scores * per cell type for each tumor across patients- **(E)** OSR; **(F)** HySR; **(G)** HSR; **(H)** UPR; and **(I)** DDR.) ifferent letters denote significant differences in stress scores as determined by ANOVA followed ’ y Tukey’s HSD test. Groups with the same letter are not significantly different from each other.

**Figure 3.**
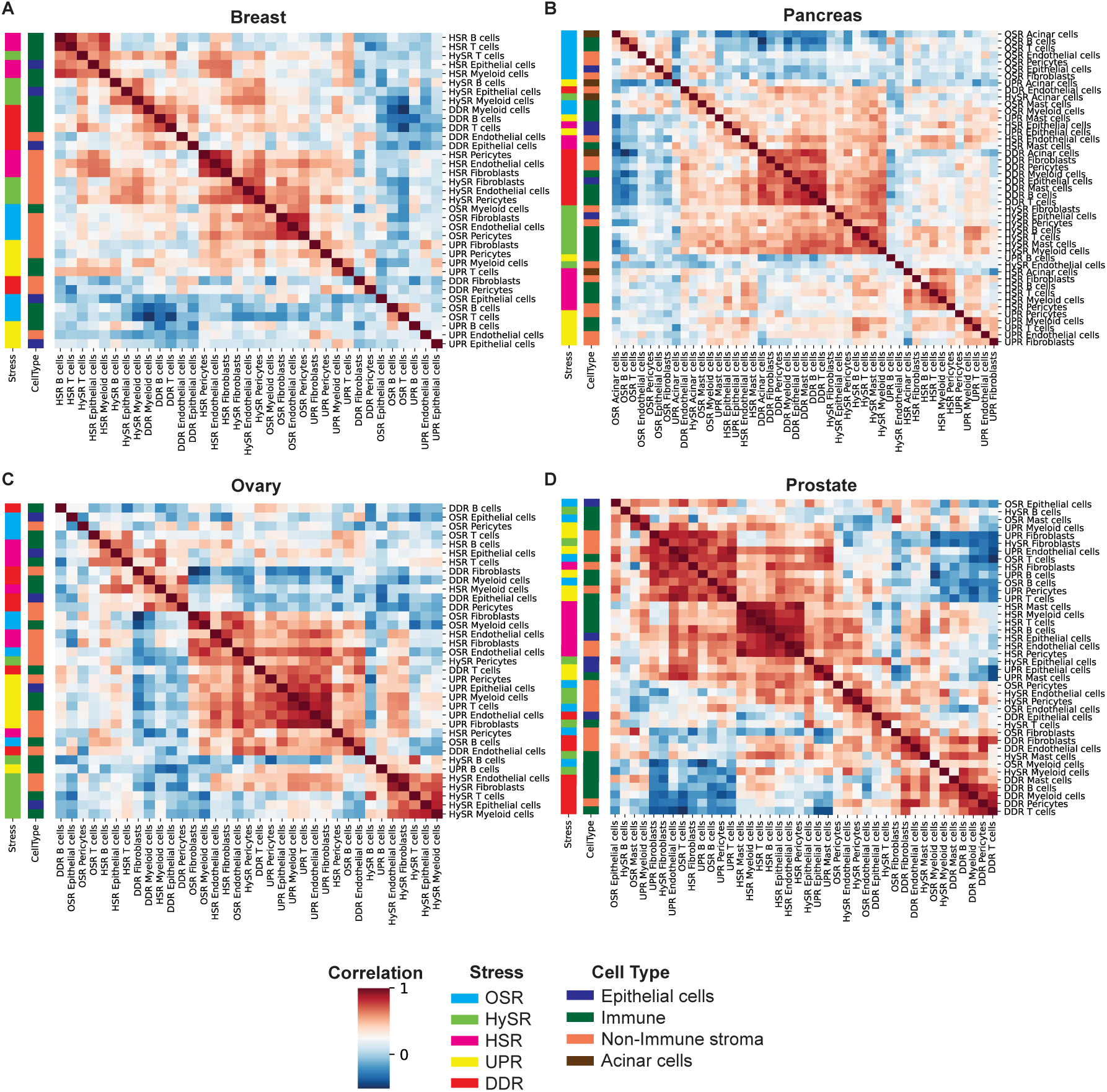
Correlation analysis of stress signatures reveals coordinated activation of the HSR in prostate tumors, the DDR in pancreatic tumors and the UPR in ovarian tumors. **(A-D)** Correlation matrix of stress scores of different cell types across patients calculated from the scRNA-seq datasets listed in Figure 2. Per-patient average scores were quantified, and Pearson coefficients of all possible pairs were calculated. Outlier patients were removed to avoid bias. Color bars indicate the stress or cell type.

### Specific subsets of CAFs and pericytes exhibit increased stress response activation

The finding that stromal cells located close to cancer cells tend to exhibit higher stress scores prompted us to test the effect of spatial positioning on the transcriptional stress signatures. To incorporate this dimension, we analyzed publicly available human cancer spatial transcriptomics data from breast, ovarian and prostate tumors (see Methods; Supplementary Figure 2A-E). We defined cell types and stress patterns using scRNA-seq data (Pal et al, 2021; Wu et al, 2021; Chen et al, 2021a; Geistlinger et al, 2021; Zhang et al, 2022; Olbrecht et al, 2021) and he stress signatures we generated, respectively. In breast and ovarian tumor slides, epithelial cells predominated, followed closely by CAFs. Roughly half of the cells in these samples belonged to the immune compartment (Supplementary Figure 2C-D). In contrast, prostate tumor slides were primarily composed of epithelial cells, CAFs, and pericytes, with a minimal presence of immune cells (Supplementary Figure 2E). We then asked which cell types are enriched within regions expressing high stress activation signatures. We found, in breast and ovarian tumors, that regions with high HSR, UPR and DDR expression were enriched with cancer/epithelial cells; while regions with high OSR and HySR were enriched with stromal cells, specifically CAFs (Supplementary Figure 2C-E). These results are consistent with the stress expression patterns we witnessed in the scRNA-seq analysis (Figure 2E-I, Supplementary Figure 1A-E), and highlight the differential stress activation between cancer and stromal cells in the TME.

While stress responses in the immune-TME were extensively studied, our understanding of the global stromal stress network is limited. Major players in the stromal microenvironment are CAFs. Recently, pericytes were also shown to contribute to the stromal TME, by transitioning into CAF-like protumorigenic cells (Sun *et al*, 2021).

CAFs were shown to divide into 3 subpopulations: myofibroblastic CAFs (myCAFs), immune-regulatory CAFs (iCAFs), and antigen-presenting CAFs (apCAFs) (Lavie *et al*, 2022). The heterogeneity of pericytes is less studied, but recent studies suggest the existence of two main subpopulations of pericytes in the TME which differ functionally (Li *et al*, 2023; Lyle *et al*, 2016). Indeed, when we re-analyzed the transcriptional landscape of fibroblasts and pericytes from the above-mentioned datasets, we found the three CAF subpopulations (Figure 4A-D; Supplementary Figure 3A-E; Supplementary Table 2). In ovarian tumors we also identified a cluster of mesothelial cells. Ovarian mesothelial cells cover the peritoneal cavity and are involved in ovarian cancer progression (Mogi *et al*, 2021). In the prostate cancer dataset, we observed a limited number of CAFs. Given that multiple studies have highlighted the presence and role of fibroblasts in prostate tumors, this could be attributed to a technical variation (Bedeschi *et al*, 2023; Bonollo *et al*, 2020). We also identified two distinct cancer-associated pericyte subpopulations, which we termed matriPer and musclePer, based on enriched pathway analysis (Figure 4A-D; Supplementary Figure 3F; Supplementary Table 3). Top upregulated pathways for matriPer were associated with the matrisome and wound healing processes, while musclePer showed an enrichment in muscle contraction pathways, indicating a distinct role for each of the pericyte subpopulations in the TME. These results align with a recent study that identified similar pericyte subsets in colorectal tumors (Li *et al*, 2023). For each fibroblast and pericyte subpopulation, we defined a unique gene signature that is shared across all four organs using differential gene expression analysis (see Methods; Supplementary Table 2, Supplementary Figure 3A-E).

**Figure 4.**
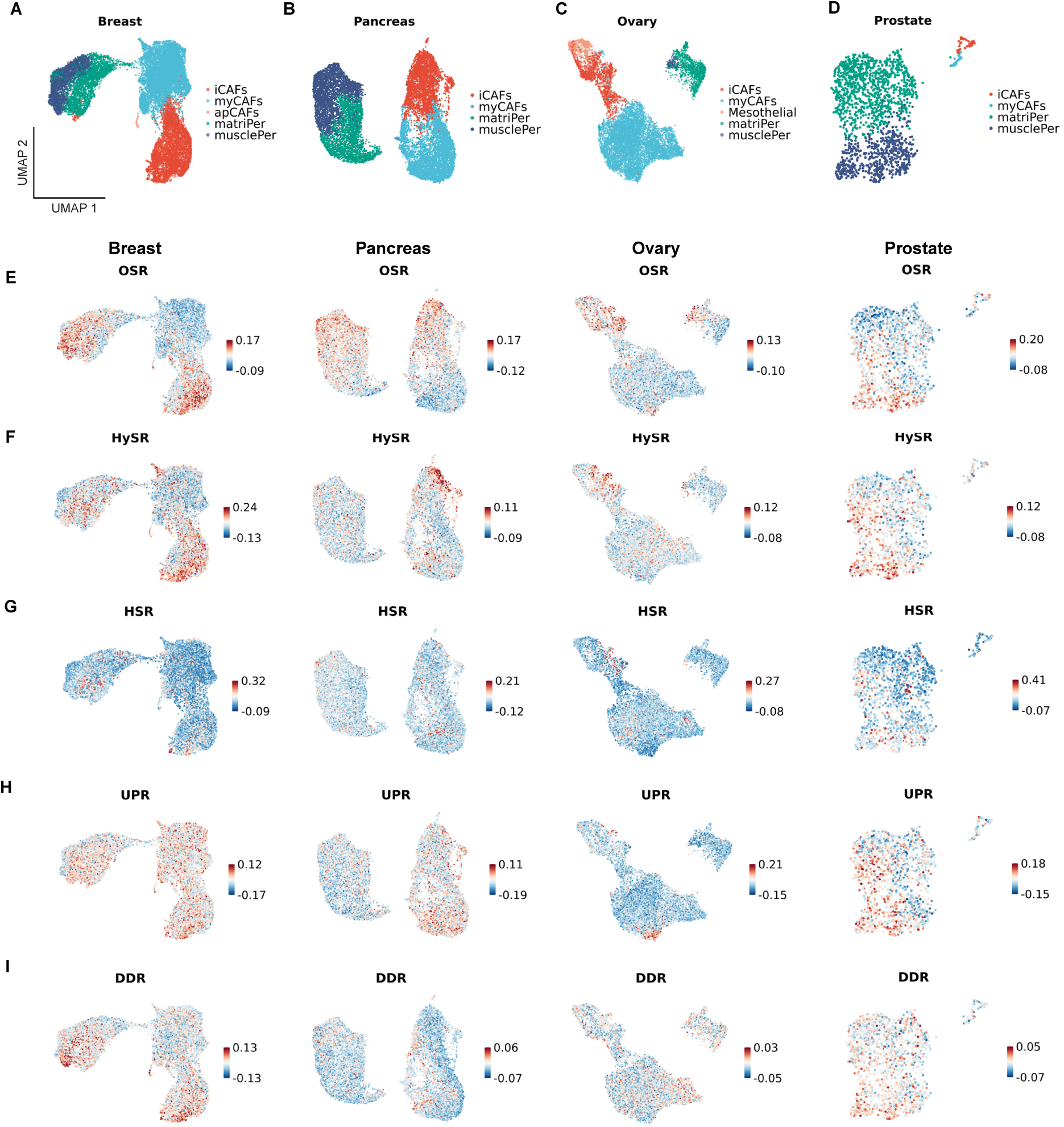
The hypoxia and oxidative stress responses are differentially activated across subpopulations of the non-immune tumor stroma. **(A-D)** UMAP plots of 20,754 fibroblasts and pericytes from 51 breast cancer patients **(A);** 14,516 fibroblasts and pericytes from 57 pancreas cancer patients **(B);** 10,762 fibroblasts and pericytes from 20 ovarian cancer patients **(C);** and 1,697 fibroblasts and pericytes from 12 prostate cancer patients **(D)**. UMAPs are colored by cell type, defined by the gene signatures we defined (Supplementary Table 2). The scRNA-seq data originates from the same datasets highlighted in Figure 2 **(E-I)** Projection of the five stress signatures scores **(E)** OSR; **(F)** HySR; **(G)** HSR; **(H)** UPR; and **(I)** DDR.

Next, to assess the stress responses activation of these cells, we projected the stress scores on the fibroblast and pericyte UMAPs (Figure 4E-I). We calculated the average scores per patient and compared them among the subpopulations of fibroblasts (Supplementary Figure 4A-E) and pericytes (Supplementary Figure 4F-J). Due to the low number of fibroblasts in the prostate dataset, we did not analyze the fibroblast subpopulations in this tumor type. Additionally, due to their vast abundance and dominant presence across the different tumors, we focused our downstream analysis on the iCAF, myCAF, MatriPer and MusclePer subpopulations. We found that the OSR score is higher in iCAFs compared to myCAFs in all three organs (Figure 4E, Supplementary Figure 4A). HySR scores were higher in breast and pancreas tumor iCAFs (Figure 4F, Supplementary Figure 4B), while DDR scores were elevated in breast iCAFs, as well (Figure 4I, Supplementary Figure 4E). Additionally, musclePer had higher OSR in all tumors, and higher HSR scores in breast, pancreas, and ovary tumors (Figure 4E,G; Supplementary Figure 4F,H). These results identify two subpopulations of the non-immune stroma - iCAF and musclePer - as stress-associated and can indicate specific roles for them in tumor progression, while phenotypically distinguishing them from other fibroblasts and pericytes.

### NRF2 and the oxidative stress response are activated in iCAFs

To further investigate the interplay between oxidative stress and stromal heterogeneity, we analyzed the correlations between stress and cell type signatures at the single cell level (Figure 5A-B; Supplementary Figure 5A-B). The iCAF and OSR signatures showed a positive correlation across all tumor types (Figure 5A), while the myCAF signature did not correlate positively with any stress response and was in fact negatively correlated with the OSR signature (Supplementary Figure 5A). In pericytes, the musclePer subpopulation showed a positive correlation with the OSR signatures (Figure 5B), while the matriPer subpopulation showed no positive correlations with any of the stress responses (Supplementary Figure 5B).

**Figure 5.**
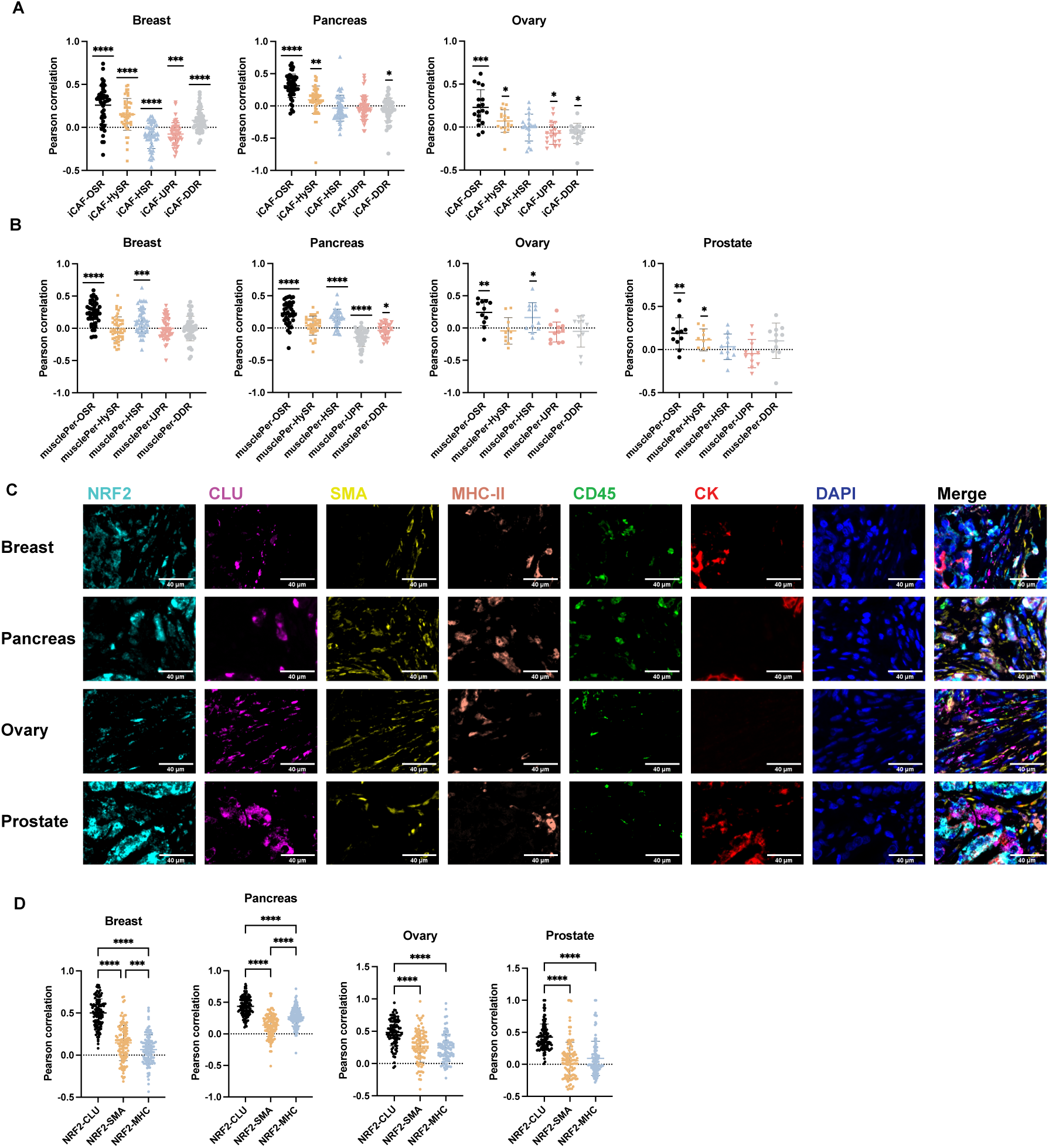
NRF2 and the oxidative stress response are associated with iCAF signature. **(A-B)** Pearson correlation coefficients between stress and cell type scores calculated from the scRNA-seq data described in Figure 4. **A** - iCAFs; **B** - musclePer. P-Values were calculated using one sample t-test (p=0). **(C-D)** Formalin-fixed paraffin-embedded (FFPE) tumor microarrays (TMAs) of breast (N = 114), pancreas (N = 125), ovary (N = 93), and prostate (N = 104) cancer patients were stained by MxIF and analyzed using QuPath software, CD45^-^ CK^-^ cells were defined as non-immune stromal cells. Representative images are shown **(C)**. Scale bar = 40pM. For each patient Pearson correlation coefficients between the staining intensities of NRF2 and the different CAF markers CLU, aSMA and MHC-II were calculated **(D)**. P-Values were calculated using one-way ANOVA, followed by Tukey’s multiple comparisons test.

To test whether these findings translate to the protein level, we stained human tissue microarrays of different tumors for markers of the three main CAF subpopulations (CLU, iCAF; aSMA, myCAF; MHC-II, apCAF) (Lavie *et al*, 2022) and for NRF2, the main regulator of the oxidative stress response, and then quantified and calculated the correlation between the intensities of NRF2 and the three CAF markers at single cell level (Figure 5C). We found that NRF2 staining positively correlated with CLU staining in the non-immune stroma of breast, pancreas, ovary, and prostate tumors. Moreover, we found that the NRF2-CLU correlation was the highest compared to all other CAF markers (Figure 5D), suggesting that the more the OSR is activated in a cell, the higher the likelihood that it is an iCAF. These findings support the conclusion that iCAFs show a stronger OSR and point to a mechanistic role for NRF2 in the regulation of the iCAF phenotype.

To test this hypothesis in an independent dataset, we analyzed patient data from the TCGA database of breast, pancreas, and ovary tumors. We aimed to assess the correlation between the different stress responses and the cellular composition of the tumor. We implemented the CIBERSORTx(Newman *et al*, 2015) algorithm to estimate the fractions of the different cell types in the tumors. We then ranked the patients from lowest to highest stress score (Figure 6A-E; Supplementary Figure 5C-L). We found that across all 3 tumor types the HySR appeared to be inversely correlated with the number of epithelial cells, and the OSR shows the same behavior in breast and ovarian tumors (Figure 6A,B; Supplementary Figure 5C-D,H-I; dark blue). The UPR, on the other hand, showed an opposite trend - the relative number of epithelial cells increased as the UPR score increased in breast cancer (Figure 6D). Additionally, for both OSR and HySR, the iCAF population (dark red) seemed to increase as the stress score increased and the epithelial cells decreased in breast and pancreas tumors (Figure 6A,B; Supplementary Figure 5C-D). These results support our claim that oxidative stress is associated with the iCAF phenotype.

**Figure 6.**
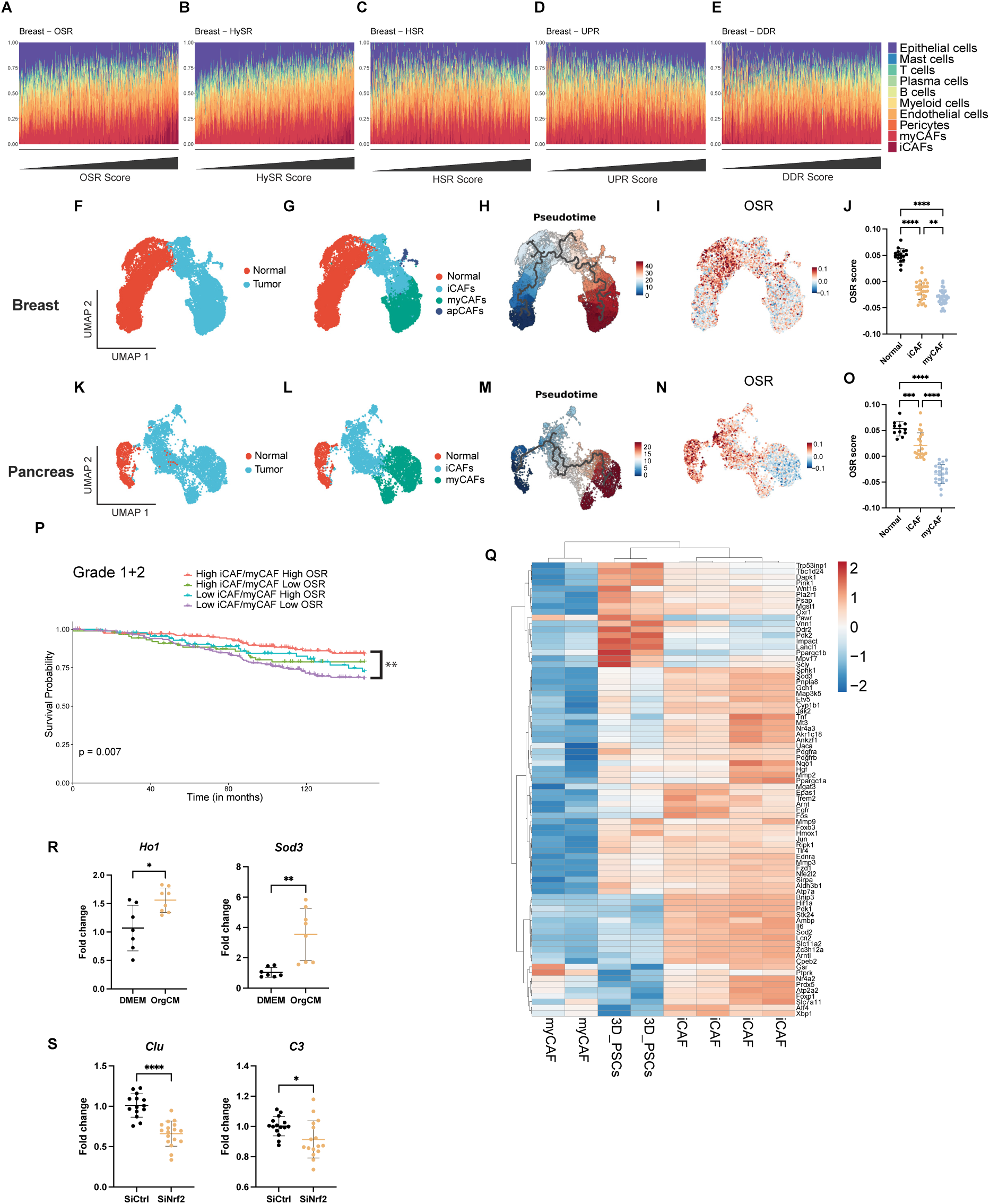
NRF2 and the oxidative stress response contribute to the transition of normal fibroblasts to iCAF. **(A-E)** Patient data from the TCGA breast cancer dataset was analyzed for cellular composition using the CIBERSORTx (Newman *et al*, 2015) algorithm and the results were ordered by the stress score. **(F-O)** Trajectory analysis. **(F,K)** UMAP plots of scRNA-seq data of breast **(E)** or pancreas **(J)** fibroblasts with data from normal samples, re-analyzed from publicly available datasets (Pal *et al*, 2021; Peng *et al*, 2019). **(G,L)** Clusters were annotated based on the stromal gene signatures we described in Supplementary Figure 3. **(H,M)** Trajectory analysis of breast and pancreas tumors and normal samples using the Monocle3 R toolkit. **(I,N)** Projections of the OSR score we previously defined. **(J,O)** Patient-level quantification of OSR scores. P-Values were calculated using one-way ANOVA, followed by Tukey’s multiple comparisons test. **(P)** Kaplan-Meier analysis of overall survival for low-grade breast cancer patients from the METABRIC cohort (Curtis *et al*, 2012). Patients were stratified based on their OSR signature and iCAF/myCAF ratio, calculated by CIBERSORTx (median was used as cutoff). P-values were calculated from the log-rank test, and paired comparisons were calculated using the Survdiff function in R with FDR correction. **(Q)** Heatmap of the differentially expressed oxidative stress related genes from bulk RNA-seq between cell culture models of iCAFs, myCAFs and quiescent PSCs, re-analyzed from (Ohlund *et al*, 2017). Differentially expressed genes were filtered by a logFC threshold of 0.5 and adjusted p-value of 0.05. **(R)** Immortalized PSCs were seeded in matrigel for 3 days with either DMEM or KPC organoid conditioned medium (orgCM) and OSR genes *Hol* and *Sod3* were measured using qPCR. **(S)** Immortalized PSCs were seeded in 2D culture and were depleted of *Nrf2* using siRNA. Cells were then seeded in Matrigel for 3 days with orgCM, and known iCAF markers in this system *Clu* and *C3* were measured using qPCR. Results are shown as mean ± SD. P-Values were calculated using two samples t-test.

Supported by our finding that NRF2 and the OSR are high in iCAFs, and since it was suggested that normal fibroblasts likely give rise to iCAFs (Houthuijzen *et al*, 2023), we hypothesized that NRF2 either regulates the transition of normal fibroblasts to iCAF, the transition from iCAFs to myCAFs, or both. To test this hypothesis, we performed trajectory analysis of two of the scRNA-seq datasets we analyzed, which also contained normal samples of breast and pancreatic tissues (Pal *et al*, 2021; Peng *et al*, 2019)(Figure 6F-J,K-O). In both breast and pancreas, pseudotime analysis revealed a gradual transition from normal fibroblasts to iCAFs and then to myCAFs (Figure 6H,M). The OSR gene signature follows an opposite trajectory - OSR is silenced as fibroblasts transition from normal fibroblasts to myCAFs (Figure 4I,N). Averaging the OSR scores per patient revealed a gradual decrease of the OSR score in the normal-iCAF-myCAF trajectory (Figure 4J,O). The pancreatic HySR was the only other stress response to show this pattern of expression (Supplementary Figure 5M-N). These results suggest a role for NRF2 and the oxidative stress response in the transition from normal fibroblasts to iCAFs and from iCAFs to myCAFs.

To assess the clinical implications of our findings, we investigated whether the levels of OSR and the relative abundance of iCAFs within the tumor were associated with patient survival. A cohort of 1053 breast cancer patients from the METABRIC dataset (Curtis *et al*, 2012) was utilized for this purpose. We used CIBERSORTx (Newman *et al*, 2015) to profile the cellular composition of the tumors. Our analysis revealed that patients with both low iCAF-to-myCAF ratios and a low OSR score showed the poorest outcome in low grade breast tumors, while patients with high iCAF-to-myCAF ratios and a high OSR score showed the best clinical outcome (Figure 6P). In high grade breast tumors, the OSR score does not appear to have a meaningful contribution to patient survival (Supplementary Figure 5O), indicating the importance of OSR in early steps of malignant progression. These results suggest that oxidative stress may be leveraged in low-grade tumors to increase the iCAF/myCAF ratio, thus improving the disease outcomes.

### NRF2 plays a role in the transition of normal fibroblasts to iCAFs

Next, we used an established cell culture model of PDAC iCAFs and myCAFs (Öhlund *et al*, 2017) to investigate the role of NRF2 and oxidative stress in the function and plasticity of the CAFs. Ohlund *et al* utilized murine pancreas stellate cells (PSCs) that were grown as either myCAFs (2D culture), quiescent PSCs (3D culture with normal growth media) or iCAFs (3D culture with cancer organoid conditioned media (OrgCM) and sequenced cells under each condition to define unique genes upregulated in each population. Using this data, we assessed the expression levels of more than 400 oxidative stress related genes. The vast majority of differentially expressed OSR-related genes was upregulated in iCAFs and quiescent PSCs compared to myCAFs (Figure 6Q). A subset of these was upregulated in quiescent PSCs compared to iCAFs, while other genes were more highly expressed in iCAFs compared to PSCs (Figure 6Q). This apparent discrepancy from our trajectory analysis could be due to changes which the PSCs undergo in cell culture, causing them to somewhat lose their normal-like phenotype.

To experimentally validate these results, we cultured PSCs in quiescent- or iCAF-inducing conditions. We confirmed their quiescent-to-iCAF transition by monitoring the expression of the iCAF genes *Clu* and *C3*, as well as the myCAF gene *Acta2*. Indeed, the iCAF genes were upregulated and *Acta2* was downregulated in the growth conditions of iCAFs (3D with orgCM) (Supplementary Figure 5P-Q). Next, to check how OSR genes are expressed in iCAFs in this model, we checked the expression of known oxidative stress genes and NRF2 targets *Ho1* and *Sod3.* We found that both genes were upregulated in the iCAF growth conditions compared to quiescent PSCs, supporting the sequencing results (Figure 6R). Finally, we silenced *Nrf2* using siRNA prior to the addition of OrgCM to PSCs in 3D culture (Supplementary Figure 5R) and measured the expression of the iCAF markers *Clu* and *C3.* We found that silencing of *Nrf2* led to downregulation of *Clu* and *C3* (Figure 5S), suggesting that NRF2 is necessary for the quiescent- PSCs-to-iCAF transition.

## Discussion

The tumor microenvironment (TME) is a complex network that consists not only of cancer cells but also a variety of other cellular players that dynamically interact with one another. Understanding the various stresses these cells undergo and the resulting cellular responses is essential for a holistic view of tumor biology and progression. Here we presented a comprehensive map of the network of stress responses in the TME, by dissecting five stress responses across four different tumor types. We found that the oxidative stress response and its central regulator NRF2 play a role in the regulation of two stromal subpopulations - iCAFs and musclePer, and showed that low-grade breast patients with high OSR and high iCAF content exhibit better survival, suggesting a protective role for OSR and iCAF. Overall, our study offers an unbiased and holistic view of the stromal stress response landscape and proves the important contribution these cellular processes have on the tumor.

We observed a spatial relationship between the level of stress responses and proximity to the tumor, suggesting non-cell-autonomous signaling within the TME. This pattern may hint at the ability of cancer cells to induce stress responses in surrounding non-malignant cells. Not only does this finding highlight the prominent effect of cancer cells on their microenvironment, it also emphasizes the dynamic interplay between cell types in the TME. This spatial gradient of stress responses could potentially be due to factors released by the cancer cells, such as cytokines, that act on the adjacent cells. This introduces the hypothesis that cancer cells may utilize these stress signals to subvert normal cell function for their benefit. For example, this gradient of stress signals could potentially influence immune cell function in the TME, enabling a more immune-suppressive and pro-tumorigenic environment (Salvagno *et al*, 2022). The spatial localization and functionality of immune cells were shown to be significantly influenced by metabolic stress (Chang *et al*, 2015). Thus, stress signals from cancer cells may serve to create an immunosuppressive microenvironment, further promoting tumor progression and resistance to therapy. Understanding the mechanisms behind this spatial gradient of stress responses could provide novel insights into tumor biology and how the cancer cells influence the non-malignant compartments of the TME. Our single-cell transcriptomic analysis revealed that while there is a certain level of universality in the stress response signatures across different tumor types, each tumor exhibits its unique pattern, suggesting an organ-specific regulation of these stress responses. This implies that stress responses in the TME are not merely reactive but could be intricate, dynamic, and tailored to the specific demands of each tumor. Whether this is driven by the mutational landscape or by organ dependencies remains to be determined and requires larger cohorts of patients. The unique stress response patterns we observed across different cell types highlight the diverse and adaptable nature of the tumor microenvironment. Endothelial cells, key components of the tumor vasculature, demonstrated the highest hypoxia score across all four tumor types. This is perhaps reflective of the poor vascularization often seen in solid tumors, which results in regions of low oxygen tension or hypoxia, a condition to which endothelial cells must adapt for survival and function (Abou Khouzam *et al*, 2021). B cells, crucial components of the adaptive immune response, exhibited the highest UPR across all tumors, possibly due to their inherent high antibody demand in response to the cancer cells (Downs-Canner *et al*, 2022). Unexpectedly, we found that immune cells, traditionally associated with immune surveillance and response, showed high levels of DDR, indicating a potential cell-non-autonomous role for the DDR (Dai *et al*, 2022). The OSR was found to be high in CAFs across all tumor types, emphasizing a potential role for the OSR in the fibroblasts. The oxidative stress experienced by these cells might contribute to their functions, including remodeling of the extracellular matrix and modulation of immune responses (Nguyen *et al*, 2018; Chan *et al*, 2017; Nicolas *et al*, 2022; Giannoni *et al*, 2011). The variation in HSR activation, particularly its high expression in breast and pancreatic cancer cells, but not as much in ovarian and prostate tumors, may hint to specific physiological or molecular differences between these tumor types.

We found the OSR to play a significant role in the behavior of CAFs and pericytes, and our findings indicate that the transcription factor NRF2, a key regulator of OSR, may be instrumental in shaping the iCAF and musclePer phenotype. We showed that musclePer cells exhibit an elevated stress response, specifically HSR and OSR and are associated with smooth muscle contraction pathways. Their exact contribution to tumor dynamics is still unexplored, as is the role of the stress responses to their functionality. Regarding iCAFs, while an upregulation of OSR genes in iCAFs was shown before in PDAC(Elyada *et al*, 2019), the matter was not pursued. This is an intriguing link, considering the importance of the OSR and NRF2 in tumorigenesis and tumor progression. We observed higher levels of OSR in iCAFs compared to myCAFs, hinting at a possible role of oxidative stress in driving the immune-regulatory phenotype of CAFs. This is further supported by the positive correlation between OSR and the iCAF signature across all analyzed tumor types. This suggests a potential role for the OSR in the development and function of iCAFs, a notion that is supported by a study that demonstrated that loss of Cav1, a known regulator of NRF2, leads to mitochondrial dysfunction, oxidative stress, and aerobic glycolysis in CAFs and induces genomic instability in adjacent cancer cells (Nguyen *et al*, 2018). This further supports our suggestion that oxidative stress in CAFs can significantly impact the development and behavior of the tumor, hinting at a possible role of oxidative stress in driving the immune-regulatory phenotype of CAFs. Further investigation is necessary to elucidate the underlying mechanisms by which OSR affects the iCAF population and the transition of the TME towards a more anti-tumorigenic state.

Toullec *et al*. showed in their study that oxidative stress can convert normal fibroblasts into (Toullec *et al*, 2010). In our study, using trajectory analysis, we found that CAFs transitioned from normal to iCAFs and then to myCAFs in two datasets of breast and pancreatic tumors, with a corresponding decrease in OSR along this trajectory. Understanding the mechanisms behind the effect of oxidative stress on fibroblasts transformations could provide valuable insights into how to leverage this process for therapeutic benefits, particularly since iCAFs have been shown to attract immune cells and enable a more anti-tumorigenic environment, compared to myCAFs (Arpinati & Scherz-Shouval, 2023).

Our analysis of the TCGA datasets of breast, ovary and pancreas tumors demonstrated a correlation between a high OSR score and an enrichment of iCAFs, alongside a reduction in epithelial cells. This suggests the OSR might be integral not only to the rewiring of CAFs but also in modulating tumor progression. Our findings suggest that a higher OSR within the TME might not necessarily promote a pro-tumorigenic environment. Instead, an elevated OSR could potentially act as a restraining factor, potentially hindering tumor growth and progression (Arfin *et al*, 2021). In the early stages of tumor development, oxidative stress and other stress responses often act as a protective mechanism, aimed at maintaining cellular integrity and preventing malignant transformation. During this phase, elevated levels of reactive oxygen species (ROS) can promote apoptosis and senescence of precancerous cells, thus serving as a defense mechanism against tumorigenesis. However, as tumors progress, they can exploit these stress responses to their advantage. Chronic and unresolvable oxidative stress can result in a dysfunctional TME, leading to genomic instability, metabolic reprogramming, and immune evasion. At this point, oxidative stress becomes pro-tumorigenic, contributing to tumor growth, invasion, and resistance to therapies. This switch from a protective to a detrimental role reflects the dual-edged sword nature of oxidative stress in cancer, highlighting the complexity of the interplay between cellular stress responses and tumor progression. Our survival analysis of the breast cancer METABRIC cohort (Curtis *et al*, 2012) supports this claim: In patients with low-grade breast tumors, a high level of oxidative stress was correlated with a favorable prognosis, suggesting that at this stage, the tumor may still be susceptible to the cytotoxic effects of ROS. The ability of cancer cells to adapt and survive under high oxidative stress might be one of the critical steps in the transition from a low-grade to a high-grade tumor. In terms of therapeutic implications, our findings suggest the potential utility of antioxidant-based therapies for low-grade tumors. However, for high-grade tumors, the effectiveness of such therapies might be limited due to oxidative stress adaptation. Instead, alternative strategies could be explored.

Cancer progression and the complex interplay within the tumor microenvironment are largely influenced by a myriad of stress responses. Our extensive analysis of these stresses reveals not just the universality of these reactions, but the unique signatures each tumor type bears. As we have uncovered, the significance of NRF2, as a central player in the OSR, emerged strongly in our study, particularly in its association with iCAFs, suggesting a complex regulatory network that modulates the TME.

## Methods

### Ethics statement

All clinical samples and data were collected following approval by Memorial Sloan Kettering Cancer Center (MSKCC; IRB, protocol #15-149) and the Weizmann Institute of Science (IRB, protocols # 186-1) Institutional Review Boards.

### Human patient samples

Human tumor microarrays (TMA) containing samples from patients were purchased from US Biomax Inc. (Figure 1: Breast - BR1503f, Pancreas - PA961f, Ovary - OV2084b, Prostate - PR807c. Figure 4: Breast - BR1191, Ovary - OV2001b, Prostate - PR1211) or assembled at MSKCC (Figure 4: Pancreas). The pancreas TMA (Figure 4) contains tumor samples from surgically resected primary pancreas ductal adenocarcinomas of patients treated at MSKCC; informed consent to study the tissue was obtained via MSK IRB protocol #15-149 and the Weizmann Institute of Science IRB, protocol # 186-1. FFPE whole tumor sections and deeply annotated demographic, clinical, pathologic and genomic (MSK-IMPACT^TM^) data were collected for all MSKCC patients in the study.

### Cell culture

Mouse-immortalized PSCs and KPC organoids were kindly provided by David Tuveson’s laboratory (CSHL, USA). PSCs were cultured in growth medium containing Dulbecco’s modified Eagle’s medium (DMEM; Biological industries, 01-052-1 A) supplemented with 10% fetal bovine serum (FBS) and 1% penicillin/streptomycin and were maintained at 37°C in 5% CO2. KPC organoids were cultured in Corning® Matrigel® Growth Factor Reduced (GFR) Basement Membrane Matrix, Phenol Red-free, LDEV-free, (Corning, 365231) with complete organoid medium (Öhlund *et al*, 2017). Conditioned medium was collected following 3-4 days of culture with 5% FBS DMEM. For 3D culture, 4*10^4^ PSCs were seeded in Matrigel® GFR in organoid conditioned media for 3 days, after which cells were harvested for further analysis.

### RNA isolation and qPCR

RNA isolation was performed using the Bio-Tri Reagent (Bio-Lab, cat. #959758027100), following the manufacturer’s instructions. Complementary DNA (cDNA) synthesis was performed with High-Capacity RNA-to-cDNA Kit (Thermo Fisher Scientific, cat. #4387406). The primer sequences used for qPCR analysis are provided in Supplementary Table 4.

### Immunofluorescent staining of the human tumor microarray

Human tumor microarrays (TMA) containing samples from patients were purchased from US Biomax Inc. (Figure 1: Breast - BR1503f, Pancreas - PA961f, Ovary - OV2084b, Prostate - PR807c. Figure 4: Breast - BR1191, Ovary - OV2001b, Prostate - PR1211. Pancreas TMAs were generously given to us by Prof. David Kelsen, MSKCC; IRB, protocol #15-149). TMAs were deparaffinized and fixed with 10% neutral buffered formalin. Antigen retrieval was performed using citrate buffer (pH 6.0) or Tris-EDTA buffer (pH 9.0). Slides were then blocked with 10% BSA + 0.05% Tween20 and the antibodies were diluted in 2% BSA in 0.05% PBST and used in a multiplexed manner with OPAL reagents (Akoya Biosciences). All primary antibodies were incubated overnight at 4 °C. Briefly, following primary antibody incubation, slides were washed with 0.05% PBST, incubated with secondary antibodies conjugated to HRP, washed again, and incubated with OPAL reagents. Slides were then washed, and antigen retrieval was performed. Then, slides were washed with PBS and stained with the next primary antibody or with DAPI at the end of the cycle. Finally, slides were mounted using Immu-mount (#9990402, Thermo Scientific). Images were taken with a Pannoramic Scan II scanner, *20/0.8 objective (3DHISTECH) or with a Phenocycler scanner (Akoya Biosciences). Images were analyzed using QuPath software (Bankhead *et al*, 2017). Cell segmentation was done using Cellpose (Stringer *et al*, 2021). Distance analysis was performed using QuPath software. In Figure 1, cancer cells and immune cells were distinguished based on morphological parameters, using QuPath. For TFs - positive staining was defined as nuclear staining. Antibodies used in this study are detailed in Supplementary Table 5.

### scRNA-seq analysis

The human single-cell RNA sequencing datasets used in this paper were analyzed using the Seurat (V4.3) pipeline (Stuart *et al*, 2019) in R v4.2.2. Cells were filtered with gene count between 200 and 5000 (6000 for the pancreas data); total molecules count smaller than 20,000 (30,000 for the pancreas data); and a mitochondrial gene percentage less than 10% for the pancreas data, 20% for the breast and prostate data and 25% for the ovary data. Cells were normalized by the Sctransform V2 method. dimensionality reduction and clustering were done using default parameters. Cell types were defined by canonical markers. For breast, pancreas, and ovary - the different datasets were integrated using Seurat v4.3. For figures 1-3, normal samples were excluded from relevant data sets. Trajectory analysis was performed using Monocle3 (Trapnell *et al*, 2014) with default parameters.

Stress signatures were derived from gene ontology database (Ashburner *et al*, 2000). Each signature comprises between 47 and 231 genes and are listed in Supplementary Table 1. Expression of stress signature genes within each signature, across the various cell types in the different tumors, can be found in Supplementary Tables 7-10. Genes included were expressed in more than 30% of at least one cell type.

Different integration methods yield different clustering results. We acknowledge that while using Harmony integration (Korsunsky *et al*, 2019) on scRNA-seq data from Peng *et al* (Peng *et al*, 2019) led to the identification of a cluster of apCAFs (Shaashua *et al*, 2022). In this study we used Seurat integration(Stuart *et al*, 2019) and did not find this cluster.

### Spatial transcriptomics analysis

Publicly available count matrices were downloaded from 10x Genomics and processed using R v4.2.0 and Seurat v4.3. Data were normalized using SCTransform V2 method. Each spot was given a prediction score for the different cell types, using publicly available scRNA-seq data as references (Pal *et al*, 2021; Chen *et al*, 2021a; Olbrecht *et al*, 2021; Geistlinger *et al*, 2021; Zhang *et al*, 2022). Analysis was performed using default parameters.

### CIBERSORTx

To estimate the fraction of the different cell types in the TCGA datasets, we used the computational deconvolution tool, CIBERSORTx, that estimates the relative abundance of individual cell types in a mixed cell population based on single cell RNA-seq profiles (Newman *et al*, 2015). CIBERSORTx results are detailed in Supplementary Table 6.

### Survival Analysis

Data was obtained from the METABRIC dataset (Curtis *et al*, 2012). Patients with missing inofrmation about tumor grade/stage were removed, as well as patients treated with chemotherapy. Patients were then stratified based on their iCAF/myCAF ratios, which were calculated by CIBERSORTx (see above), and based on the OSR scores. Kaplan Meier (KM) analysis of overall survival with log rank p value was performed on patients stratified by median expression of each of these signatures.

### Statistical analysis

Statistical analysis and visualization were performed using R 4.2.2, and Prism 9.2.0 (Graphpad, USA). Statistical tests were performed as described in each Figure legend. * p < 0.05, ** p < 0.005, *** p < 0.0005.

## Supplementary Figures

**Supplementary Figure 1.**
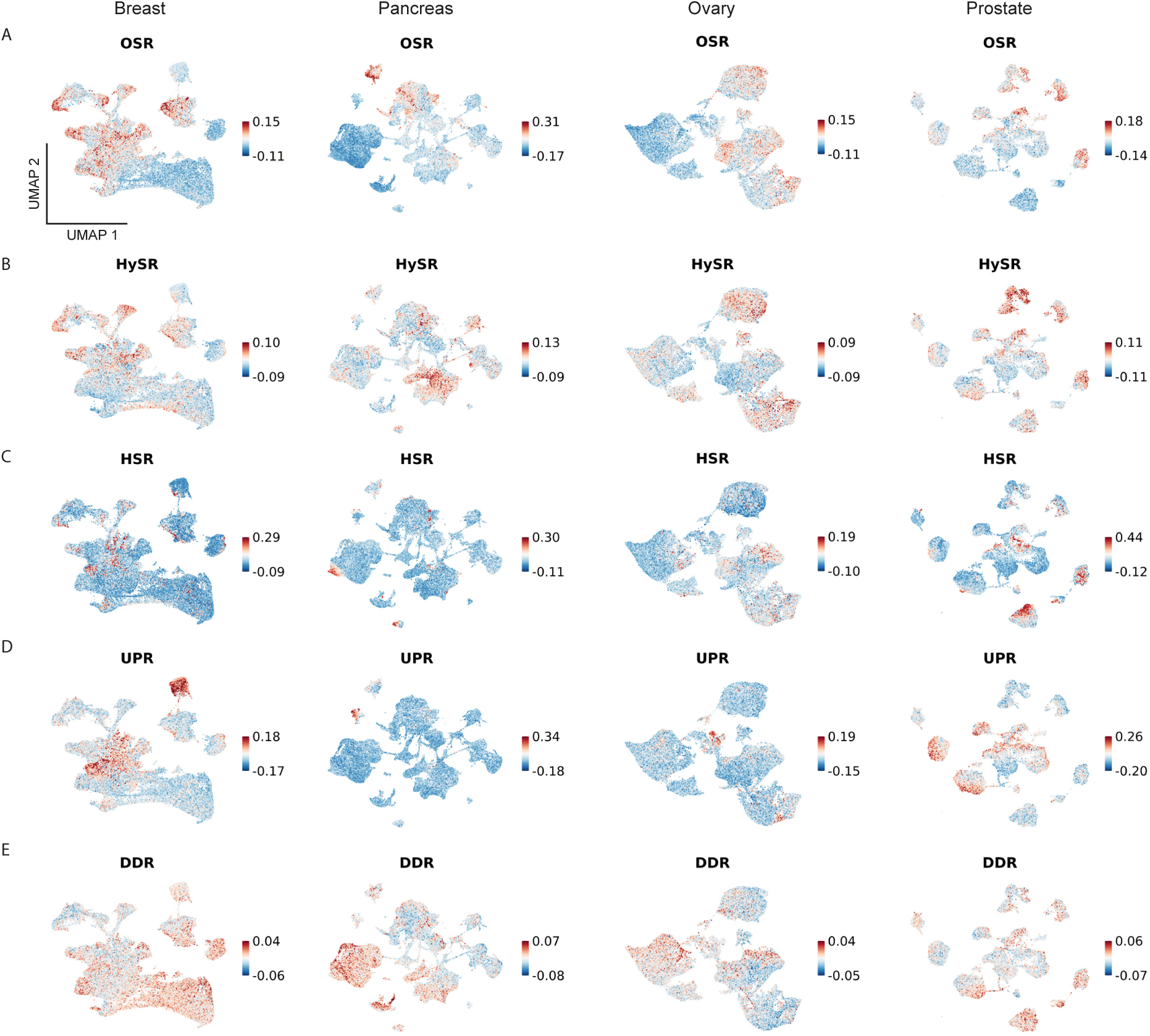
scRNA-seq data analysis uncovers shared and unique stress response activation patterns across different tumors. **(A-E)** scRNA-seq data from human tumors was reanalyzed using the Seurat R toolkit. Stress signatures (Supplementary Table 1) were projected on scRNA UMAP plots. **(A)** OSR; **(B)** HySR; **(C)** HSR; **(D)** UPR; and **(E)** DDR.

**Supplementary Figure 2.**
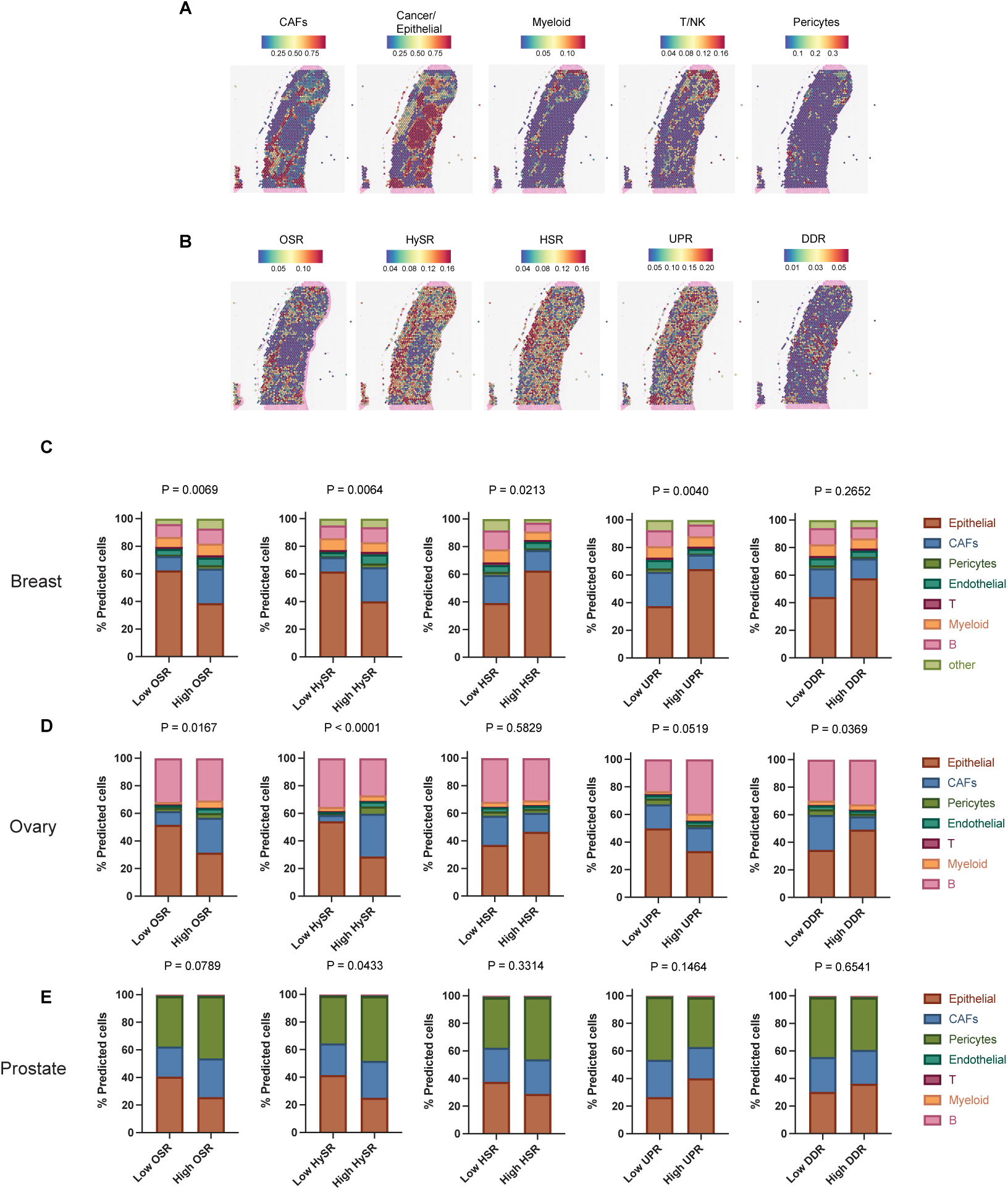
Spatial transcriptomic analysis oxidative stress signature in CAFs. **(A-B)** Publicly available human cancer slides from the 10X genomic website (www.10xgenomics.com/resources/datasets) were re-analyzed using the Seurat R toolkit. Slides from 3 breast, 2 ovarian, and 3 prostate tumors were analyzed. A slide from one breast cancer patient is presented. Using our analysis of scRNA-seq data we deconvoluted the Visium spatial transcriptomic data to predict cell type distribution in each Visium spot **(A)**. The stress signatures we defined were projected on the spatial transcriptomics data **(B)**. **(C-E)** Quantification of each tumor type. Per patient, we defined each Visium spot as expression either low or high stress for each of the stress responses (stratification based on mean), and then averaged the predicted percentage of the various cell types. **(C)** - breast, **(D)** - ovary, **(E)** - prostate. P Values were calculated using Chi-test.

**Supplementary Figure 3.**
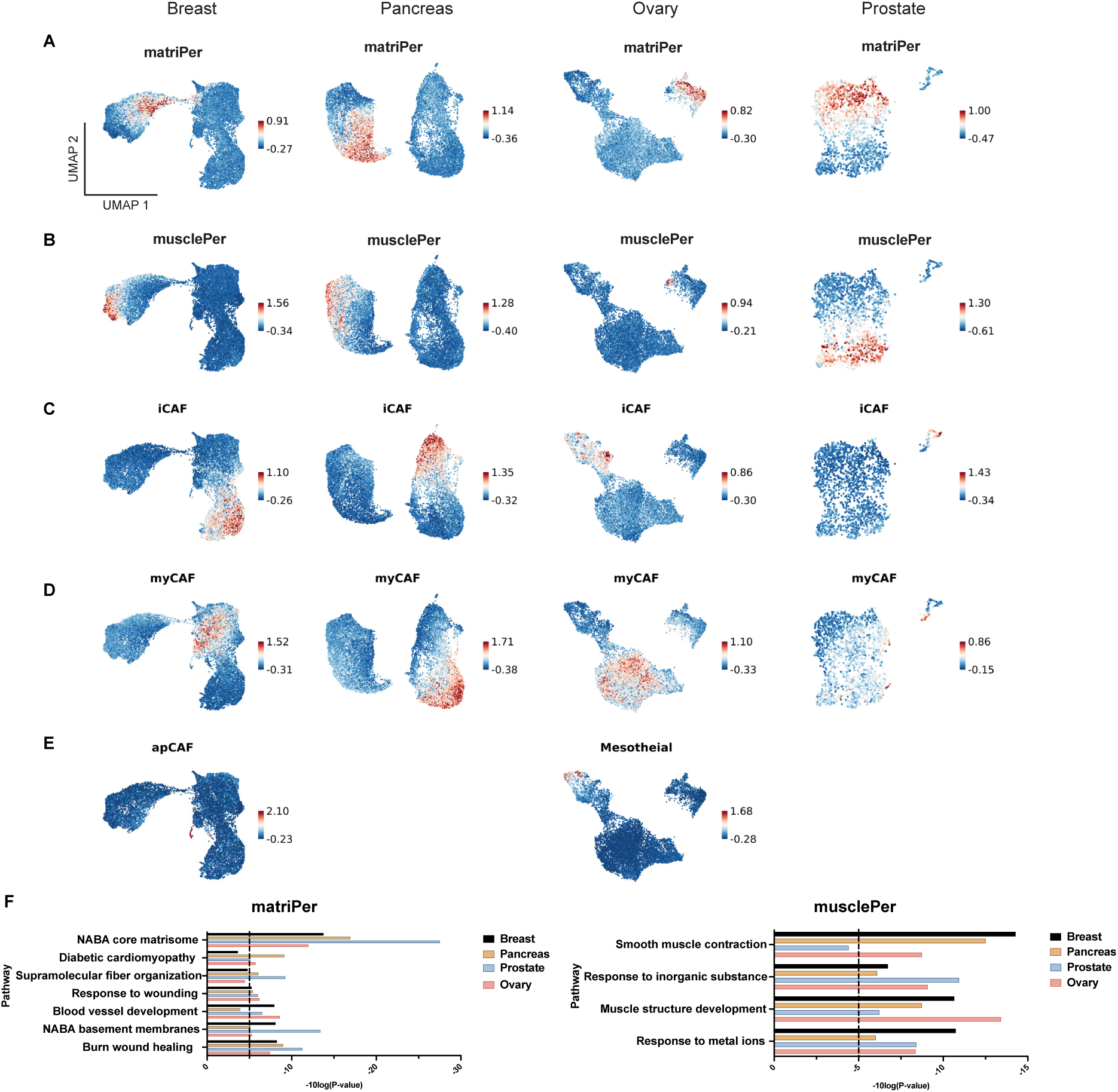
Cancer-associated pericytes are composed of two distinct subpopulations. Pan-cancer fibroblast and pericyte subpopulation signatures were defined using differential gene expression analysis of the four datasets. **(A-E)** Signature projections of the subtype signatures, which are presented in Supplementary Table 2. **(F)** Pathway analysis of two distinct cancer associated pericyte subpopulations.

**Supplementary Figure 4.**
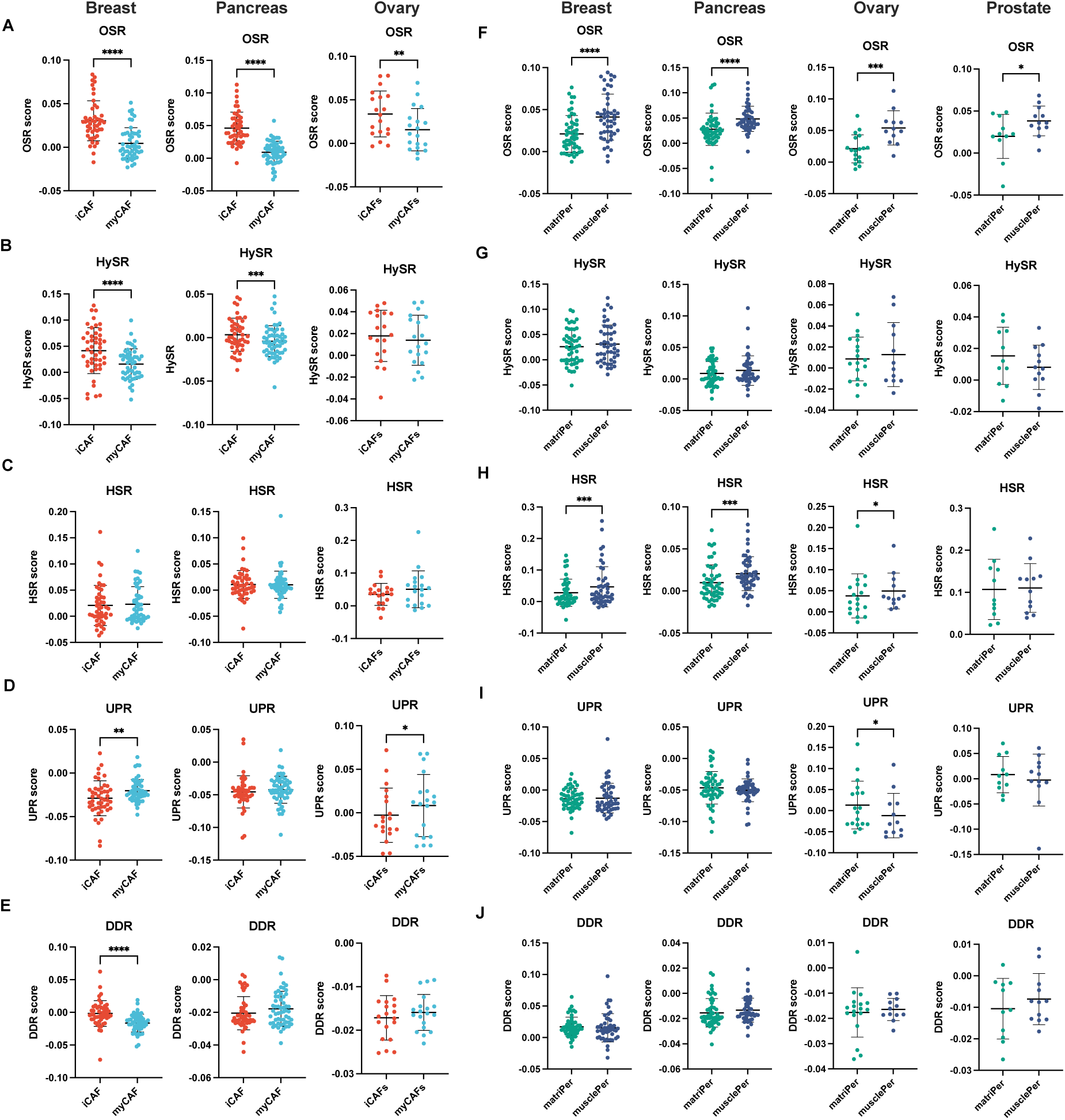
scRNA-seq analysis highlights distinctive activation of hypoxia and oxidative stress responses in various subpopulations of the tumor non-immune stromal cells. Single-cell RNA-seq data from human tumors was reanalyzed using the Seurat R toolkit. Quantification of stress scores in the different fibroblast **(A-E)** and pericyte **(F-J)** subpopulations of each tumor type, as presented in Figure 4. **(A,J)** OSR; **(B,G)** HySR; **(C,H)** HSR; **(D,I)** UPR; and **(E,J)** DDR. P-Values were calculated using the paired two-sided Student t-test.

**Supplementary Figure 5.**
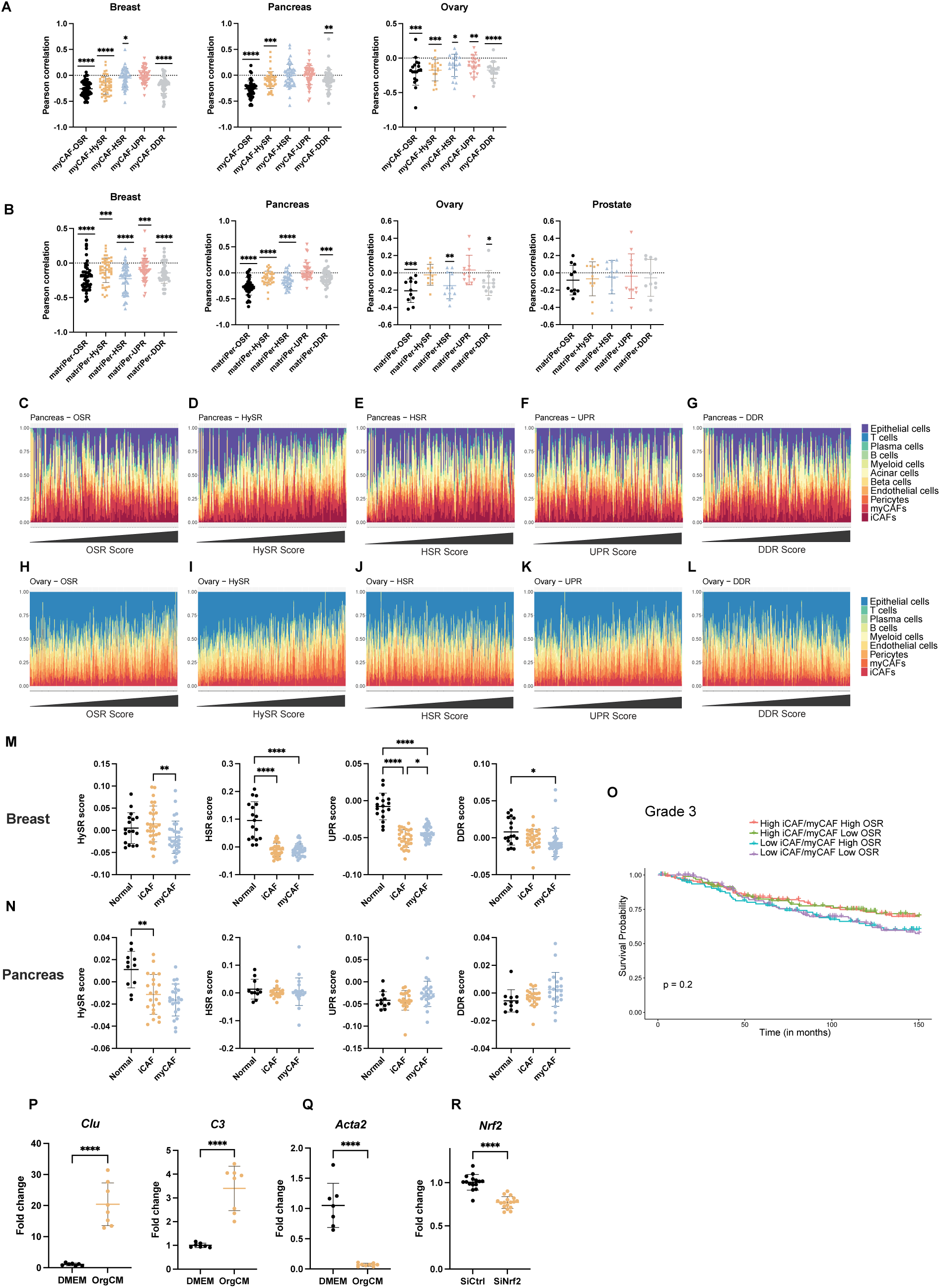
The relative amount of epithelial cells is negatively correlated with OSR and HySR scores in pancreas and ovary tumors. **(A-B)** Pearson correlation coefficients between stress and cell type scores in breast and pancreas tumors, calculated from the scRNA-seq data. **A** - myCAFs; **B** - matriPer. P-Values were calculated using one sample t- test (p=0). **(C-L)** Patient data from the TCGA pancreas **(C-G)** and ovary **(H-L)** cancer datasets were analyzed for cellular composition using the CIBERSORTx (Newman *et al*, 2015) algorithm and the results were ordered by the stress score. **(M-N)** patient-level quantification of stress scores of breast **(D)** and pancreas **(E)** tumors, calculated from scRNA-seq data of normal and tumor samples (Pal *et al*, 2021; Peng *et al*, 2019). P-Values were calculated using one-way ANOVA, followed by Tukey’s multiple comparisons test. **(P)** Kaplan-Meier curves of overall survival for high-grade breast cancer patients from the METABRIC cohort (Curtis *et al*, 2012). Patients were stratified based on their OSR signature and iCAF/myCAF ratio, calculated by CIBERSORTx (median was used as cutoff). P-values were calculated from the log-rank test and paired comparisons were calculated using the Survdiff function in R with FDR correction. **(P)** Immortalized PSCs were seeded in matrigel for 3 days with either DMEM or KPC organoid conditioned medium (orgCM) and iCAF **(P)** and myCAF **(Q)** genes were measured using qPCR. **(R)** Immortalized PSCs were seeded in 2D culture and were depleted of *Nrf2* using siRNA, and knockdown was measured using qPCR. Results are shown as mean ± SD. P-Values were calculated using two samples t-test.

